# Rosin Soap Exhibits Virucidal Activity

**DOI:** 10.1101/2021.07.19.452918

**Authors:** Stephen H Bell, Derek J Fairley, Hannele Kettunen, Juhani Vuorenmaa, Juha Orte, Connor G G Bamford, John W McGrath

## Abstract

Chemical methods of virus inactivation are used routinely to prevent viral transmission in both a personal hygiene capacity but also in at-risk environments like hospitals. Several ‘virucidal’ products exist, including hand soaps, gels and surface disinfectants. Resin acids, which can be derived from Tall oil produced from trees, have been shown to exhibit anti-bacterial activity. However, whether these products or their derivatives have virucidal activity is unknown. Here, we assessed the capacity of Rosin soap to inactivate a panel of pathogenic mammalian viruses in vitro. We show that Rosin soap can inactivate the human enveloped viruses: influenza A virus (IAV), respiratory syncytial virus and severe acute respiratory syndrome coronavirus 2 (SARS-CoV-2). For IAV, rosin soap could provide a 100,000-fold reduction in infectivity. However, Rosin soap failed to affect the non-enveloped encephalomyocarditis virus (EMCV). The inhibitory effect of Rosin soap against IAV infectivity was dependent on its concentration but not dependent on incubation time nor temperature. Together, we demonstrate a novel chemical inactivation method against enveloped viruses, which could be of use in preventing virus infections in certain settings.

**Importance:** Viruses remain a significant cause of human disease and death, most notably illustrated through the current Covid-19 pandemic. Control of virus infection continues to pose a significant global health challenge to the human population. Viruses can spread through multiple routes, including via environmental and surface contamination where viruses can remain infectious for days. Methods to inactivate viruses on such surfaces may help mitigate infection. Here we present evidence identifying a novel ‘virucidal’ product in Rosin soap, which is produced from Tall oil from coniferous trees. Rosin soap was able to rapidly and potently inactivate influenza virus and other enveloped viruses.

## Introduction

Even before the current pandemic of SARS-CoV-2, the virus that causes coronavirus disease 19 (Covid-19), respiratory-borne viruses were a leading cause of global morbidity and mortality (Institute for Health Metrics and Evaluation 2019). By way of an example, viruses such as influenza viruses (which includes IAV), are responsible for hundreds of thousands of deaths annually (Iuliano et al., 2018). To date, the pandemic of SARS-CoV-2 claimed the lives of over 3.5 million people and >180 million cases have been reported worldwide (World Health Organisation (WHO), 2021). Strategies to treat and control the spread of viruses, such as antiviral therapies and vaccines, are employed to protect the health and well-being of the general population in particular for those in at-risk settings, such as in hospital care and in the care sector (Kanamori et al., 2020). Pathogenic respiratory viruses may spread directly from person-to-person via small droplets or aerosols as well as direct contact with each other and from contaminated surfaces or fomites (Leung, 2021). Furthermore, aerosolization of environmentally-contaminated infectious virus has been observed and can spread disease (Asadi et al., 2020 and Greenhalgh et al., 2021). Infectious SARS-CoV-2 has been shown to persist on surfaces such as metal and plastic for up to 3 days respectively (van Doremalen et al., 2020). An additional layer of defence against infectious agents like viruses is the destruction of their survival on surfaces.

The infectious particle of many respiratory viruses is encased in a phospholipid bilayer or ‘envelope’, which is essential for infectivity (Cohen, 2016). For infection, enveloped viruses fuse their lipid envelope with the outer membrane, whether that is the plasma membrane or from vesicles, of the target host cell. Enveloped viruses include but are not limited to: influenza viruses, CoVs, paramyxoviruses and pneumonviruses. By comparison, non-enveloped viruses include adenoviruses and picornaviruses, such as rhinovirus and EMCV. A range of virus inactivation methods exist that can reduce the likelihood of survival or transmission of viruses via direct contact or fomites by disrupting the lipid membrane of enveloped viruses (Chaudhary et al., 2020). Such virucidal products include those targeted for personal hygiene use such as soaps or hand-gels that can be targeted to high-touch surfaces like the hands (Chaudhary et al., 2020; Chin et al., 2020). Additional measures are those that target the environment, such as surface disinfectants (Rabenau et al., 2005; Fathizadeh et al., 2020; WHO, 2020).

During a pandemic there is likely to be an increased demand for products that eliminate viral infectivity from surfaces. Coniferous trees and some other plants produce liquid resin which seals wounds in tree bark and protects the plant against pathogens and herbivores. Coniferous rosin contains resin acids, such as abietic and dehydroabietic acid, which are lipid-soluble diterpenoid carboxylic acids (San Feliciano et al., 1993). Resin acids have been shown to have antibacterial properties especially against Gram-positive bacteria (Söderberg et al., 1990; Savluchinske-Feio et al. 2006). Rosin can be collected from naturally occurring trees, but a commercially more important source of resin acids is crude tall oil, a side-stream of the cellulose processing industry. Here, we aimed to determine whether Rosin soap exhibited virucidal activity against clinically relevant pathogenic human viruses. Viruses used in this study include the enveloped viruses, IAV, RSV, SARS-CoV-2 and EMCV. Initially, 2.5% rosin soap was evaluated for its virucidal activity for all enveloped viruses examined using liquid phase assays under standardised laboratory conditions. Here, we demonstrate the potent virucidal activity of rosin soap against pathogenic enveloped viruses, supporting its further development as a surface disinfectant.

## Materials and Methods

### Cell culture

Mammalian cell lines (MDCK; Madin-Darby Canine Kidney, and Vero African Green Monkey cells) were cultured in DMEM (high glucose) supplemented with foetal bovine serum (v/v 5%) and penicillin/streptomycin (v/v 1%). Cell cultures were maintained in flasks (T175cm^2^) and passaged routinely.

### Viruses

Stocks of representative viral strains, including influenza A virus (Udorn, WSN), respiratory syncytial virus (RSV-A2), SARS-CoV-2, and encephalomyocarditis virus (EMCV)) were prepared using standard virology techniques on Vero (SARS-CoV-2 and EMCV), MDCK (influenza A virus) and Hep-2 (RSV) cells. For culture of IAV-Udorn, serum free media was used supplemented with TPCK-treated trypsin (Sigma Aldrich) at a concentration of 1ug/ml. Infectious stocks were produced and titrated in their respective cell lines before use in virucidal activity experiments. All virus work was carried out in the Biological Safety Level (BSL) 2 or BSL3 (SARS-CoV-2) facilities at QUB.

### Tall oil

The Rosin soap was produced from crude Tall Oil by Forchem Ltd (Rauma, Finland). It was a water solution obtained from dried Rosin salt consisting less than 10% sodium salts of Tall Oil fatty acids and over 90% sodium salts of resin acids. The resin acids and fatty acids of the product originated from the coniferous trees *Pinus sylvestris L.* and *Picea abies L.* The most abundant resin acid types include abietic acid, dehydroabietic acid, pimaric acid and palustris acid.

### Inactivation protocol

Virus inactivation assays were carried out in 96 well plates. Initially complete DMEM (100 μl) was added to each well, except the first column, which was used to incubate virus and product. Three concentrations of rosin soap powder were tested in each condition in duplicate (2.5%, 0.25% and 0.025% w/v). Rosin soap (Forchem Ltd (Rauma, Finland)) was dissolved in ddH_2_O. The negative control contains no virus and was incubated at 37°C. To each well of the first column, 100 μl of treatment and 100ul of virus was added. After exposure to experimental conditions, which were time (5, 15 or 10 min) and temperature (4°C, room temperature and 37°C), tenfold serial dilutions were carried out. Following dilution of the virus, permissive cells were added (100 μl) and incubated for 2-3 days. Viral infectivity was measured as the reciprocal of the final dilution giving cytopathic effect following manual investigation with a light microscope.

### Filtration

To remove residual Rosin soap from the treated virus inoculum and hence lower the level of cytotoxicity of the treatment when measuring infectivity of virus preparations, Amicon® Ultra-15 Centrifugal Filter Units (Merck) were used. 100 μl of virus (WSN) was added to 100μl of Rosin soap (2.5%) for 5 minutes at room temperature. The 200μl (WSN/Rosin Soap Powder) was washed through the filter units four times with 12ml fresh DMEM (supplemented with fetal bovine serum (v/v 5%) and penicillin/streptomycin (v/v 1%). 10-fold serial dilutions are carried out. Following dilution of the virus, permissive cells were added (100μl) and incubated for between 2-3 days. Viral infectivity was measured as the reciprocal of the final dilution giving cytopathic effect.

## Results

### Rosin Soap reduces the infectivity of influenza virus

Novel solutions to disrupt the transmission of human pathogens are required. Given that they exhibit antibacterial activity we hypothesized that Rosin soap may inhibit the transmission of pathogenic human viruses by killing viruses on surfaces. We thus took Rosin Soap and assessed whether it could reduce the infectivity of IAV (WSN strain). We chose IAV because it is a model enveloped human RNA virus and is a significant human pathogen. Furthermore, IAV achieves high titres during propagation in cell culture and is highly cytopathic (rapid cell death and rounding) in traditionally used cell lines such as MDCK cells, which together allow the facile and sensitivity determination of high levels of inhibition to viral infectivity. To determine whether Rosin Soap Powder could reduce the infectivity of IAV, we incubated influenza virus stocks with rosin acid (2.5% w/v) at 37°C for 30 minutes and measured residual infectivity by limiting dilution and assessment of cytopathic effect 72hrs later, in comparison to virus and DMEM only controls respectively. In these initial experiments, incubation of IAV with Rosin Soap Powder gave at least a ten-thousand-fold reduction in infectivity (**Fig 1**).

**Fig 1.**
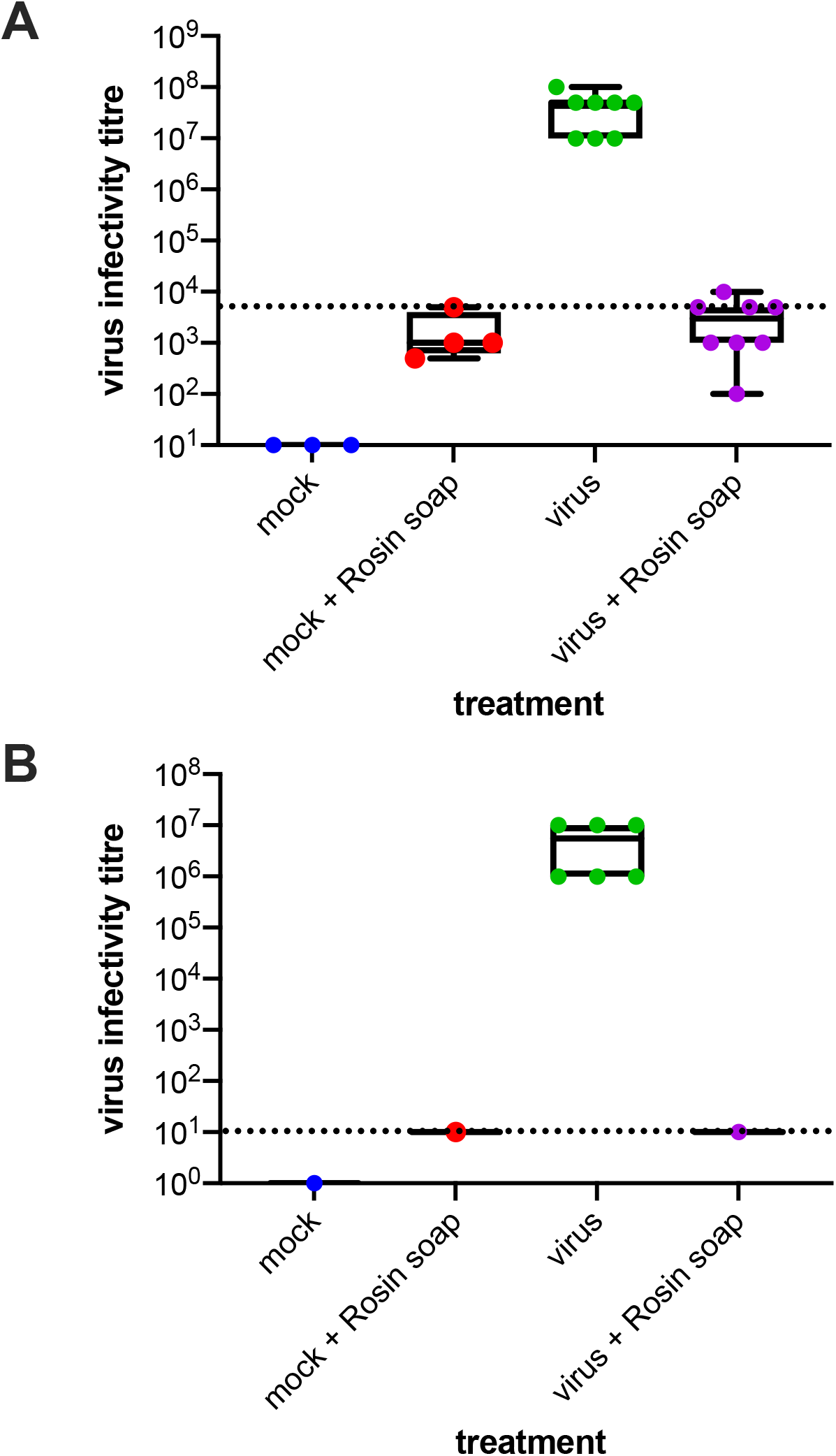
Effect of Rosin soap treatment on IAV (WSN strain) infectivity in solution compared to mock (DMEM) without (A) and with removal of residual Rosin Acid by filtration (B). IAV suspension was incubated with Rosin soap solution at 37°C for 5 minutes before residual infectivity was determined via dilution on susceptible cells (MDCK cells). Infectious virus titre corresponds to the reciprocal of the final dilution giving virus-induced cytopathic effect. Background (dashed lines) delineates the dilution that the Rosin soap treatment was toxic to the MDCK cells.

Precluding a precise determination of reduction in infectivity is the relatively high limit of detection in this assay due to the cytotoxic effect of residual Rosin Soap on cell viability, which is necessary for detection of IAV infectivity. Throughout our studies we were hindered by the relatively high cytotoxicity of rosin acids at the maximum concentration on the cells used to measure residual viral infection. This relatively high limit of detection prohibited us from determining whether there existed any viral infectivity remaining. To decrease the cytopathic effect and thus reduce the background, we filter purified our virus/rosin acid preparations prior to infectivity measurements. Experimental conditions were room temperature for 5 minutes. These experiments demonstrated a removal of the background cytotoxicity and lower limit of detection: Enhanced virucidal activity against IAV (1000000-fold) was observed (i.e only 0.00001% remaining). These data suggest that Rosin soap very likely can inactivate all infectious virus particles in each sample at 2.5%, although we cannot formally prove this.

### Assessment of the virucidal breadth of Rosin Soap

Given its effect on IAV infectivity, we hypothesized that Rosin Soap may also inhibit other viruses. To this end we investigated the virucidal activity of rosin soap against another IAV strain (H3N2, Udorn), RSV and SARS-CoV-2, as well as the non-enveloped encephalomyocarditis virus (EMCV). RSV and SARS-CoV-2 are representatives from two groups of viruses, the pneumoviruses and the coronaviruses and are themselves significant human pathogens. EMCV is a model non-enveloped virus and a pathogen of pigs and other mammals, such as non-human primates. We carried out the same protocol as above used for IAV WSN and measured residual viral infectivity using virus specific-specific means. Conditions for these experiments were room temperature for 5 minutes. In this series of experiments, all enveloped viruses were inhibited by Rosin Soap although to different degrees demonstrating that the activity of rosin soap is not limited to WSN nor IAV (**Fig 2**). In all cases, treatment with Rosin Soap brought infectivity down to baseline and fold inactivation was thus highly dependent on the starting concentration (e.g. greatest for Udorn and lowest for SARS-CoV-2). However, essentially all infectivity was brought to below the limit of detection, which is highly suggestive of near-complete inhibition of infectivity (see previous experiment). Interestingly, neither rosin soap nor Triton X (data not shown) inhibited the non-enveloped EMCV. The susceptibility of enveloped viruses to Rosin acids (and not the non-enveloped virus) suggests that the viral lipid membrane is a major target of inactivation.

**Fig 2.**
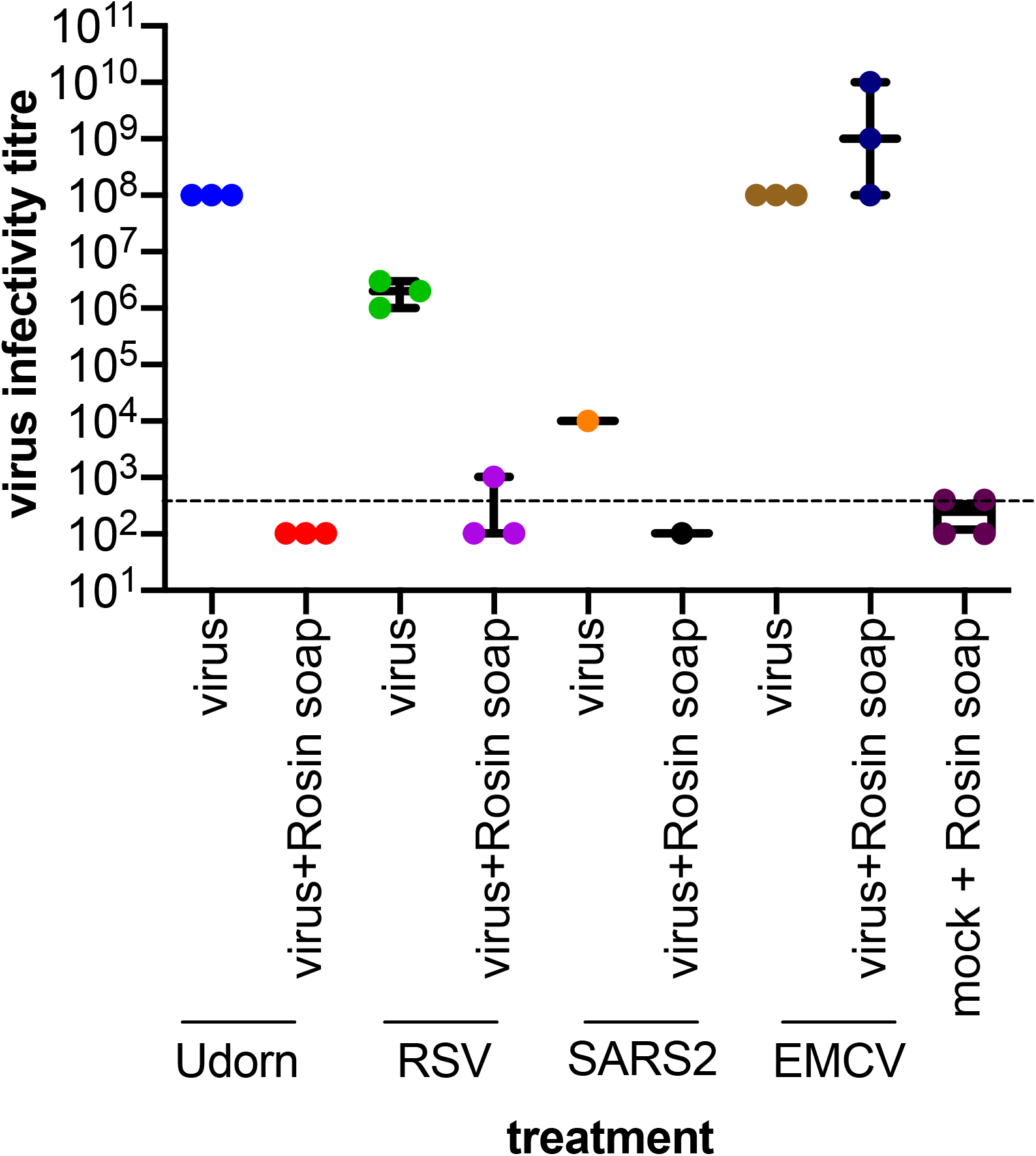
Effect of Rosin soap treatment on a panel of virus infectivity in solution compared to mock (DMEM). Three enveloped (IAV Udorn strain; RSV and SARS-CoV-2 [SARS2]) and one non-enveloped (EMCV) virus were used. Virus suspensions were incubated with Rosin Acid solution at 37 °C for 5 minutes before residual infectivity was determined via dilution on susceptible cells (MDCK cells for IAV, Vero cells for RSV, S2 and EMCV). Infectious virus titre corresponds to the reciprocal of the final dilution giving virus-induced cytopathic effect. Background (dashed lines) delineates the dilution that the Rosin soap treatment was toxic to the susceptible cells.

### Virucidal activity of Rosin soap is dependent on concentration

To understand more about the physiochemical dependence of rosin soap exhibited potent activity against enveloped viruses like IAV, RSV and SARS-CoV-2, we next determined the effect of Rosin Soap concentration, temperature and incubation time on its virucidal activity. All previous experiments were carried out with a concentration of 2.5% (w/v), time of 5 minutes and at room temperature so here we decided to alter the concentration (2.5, 0.25 and 0.025%) together with incubation time (5, 15 and 30 mins) and incubation temperature (37 °C, room temperature or 4 °C). Across all experiments, virucidal activity of Rosin Soap was only dependent on the concentration, with 2.5% showing seemingly complete activity against IAV and reduction in inhibition observed for each reduction in concentration (**Fig 3A**). In contrast to concentration, virucidal activity was independent of incubation temperature (4, room temperature [RT] or 37 °C) and incubation time with there being little difference between a 5-minute incubation compared to a 30-minute incubation (**Fig 3A-C**). These data demonstrate the rapid and efficacious activity of Rosin Acids against the enveloped virus IAV only when a critical concentration threshold has been reached.

**Fig 3.**
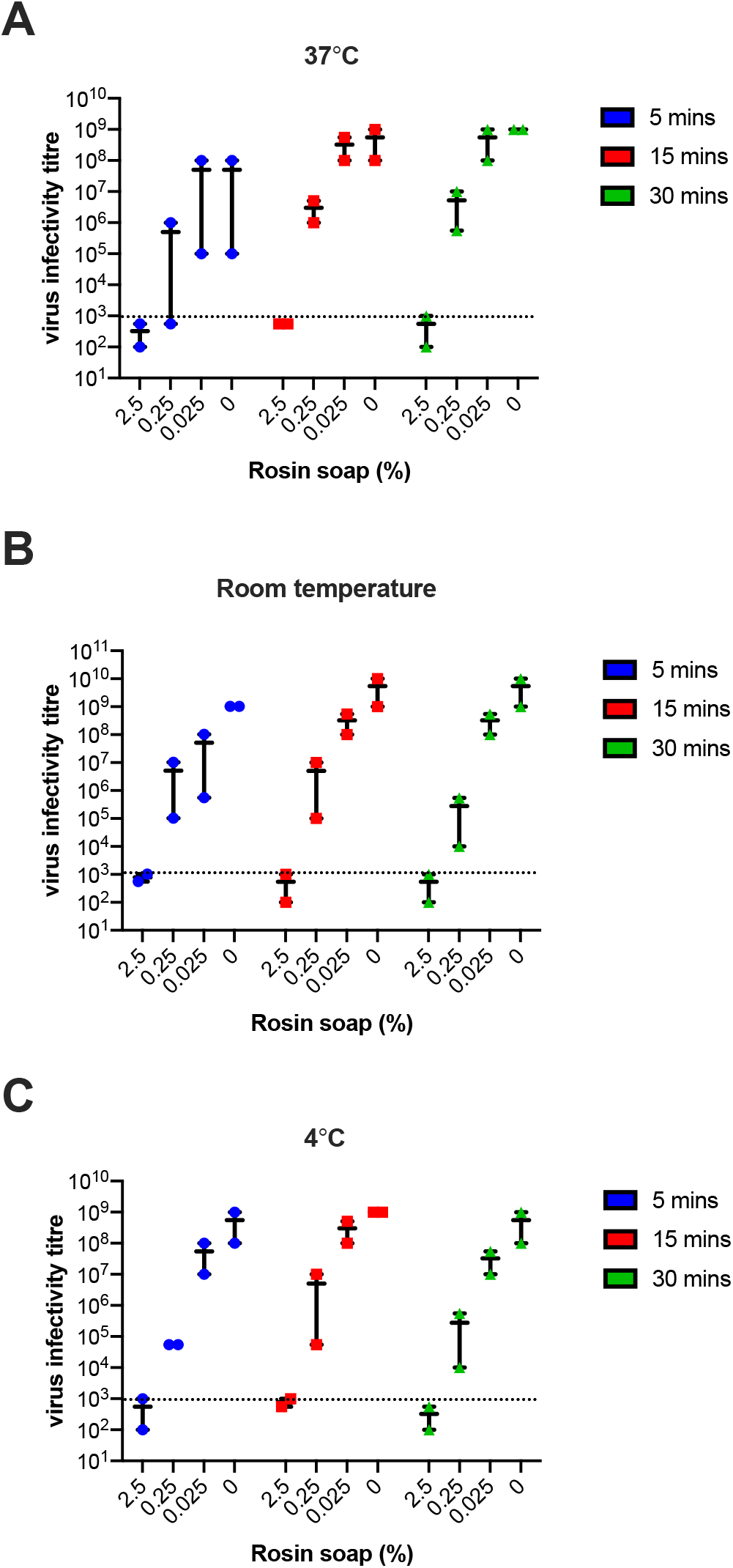
Effect of Rosin soap treatment on a IAV infectivity in solution compared to mock (DMEM) at different concentrations, treatment times and temperatures. IAV suspensions were incubated with Rosin soap solution (final concentration: 2.5, 0.25 or 0.025%) under distinct conditions before residual infectivity was determined via dilution on susceptible cells (MDCK). The effect of temperature: 37 °C (A), room temperature (B) or 4 °C (C) is shown alongside incubation time: 5 (blue), 15 (red) and 30 minutes (green). Infectious virus titre corresponds to the reciprocal of the final dilution giving virus-induced cytopathic effect. These experiments were carried out in three replicates in two independent experiments.

## Discussion

Infectious pathogenic human viruses, including SARS-CoV-2, can persist in the environment for extended periods of time, facilitating transmission via direct contact and/or through environment contamination (Marquès et al., 2020). Strategies to eliminate such infectivity from such inanimate and animate surfaces is required. Exporation of strategies that are of natural origin are warranted. To this end we sought to investigate whether Rosin soap has antiviral activity due to its reported antibacterial activity (Söderberg et al., 1990). Our work presented here shows that Rosin soap also exhibited rapid and potent viricidal activity against pathogenic human enveloped viruses but was not effective against a prototypic non-enveloped virus, EMCV.

Critically, we investigated the limits of Rosin soap virucidal activity by altering temperature, concentration and time of incubation. Using IAV as a model enveloped virus, this virucidal effect was dependent on concentration of product rather than incubation temperature or time. Interestingly, while viricidal activity of Rosin soap was not influenced by length of exposure (5 to 30 minutes) or incubation temperature (4°C, room temperature and 37°C), the only factor that did influence viricidal activity was Rosin soap concentrations with 2.5% (w/v) being the most effective. Higher concentrations of Rosin soap led to the rapid and potent loss of infectivity of IAV and other enveloped viruses. The fact that temperature nor time had a major impact of efficacy suggests that this product has highly potent virucidal activity.

Mechanistically, our results showing the lack of efficacy against non-enveloped viruses suggests that the target for Rosin soap antiviral activity is the viral envelope, which is composed of a phospholipid bilayer. The virus envelope is critically required for infectivity facilitating protection of genomic material and facile entry (catalysed by viral fusion protein machinery) into target host cells via virion-to-cell membrane fusion either at the plasma membrane or endosomal compartment membranes (Dimitrov 2004). Loss of virion envelope integrity will prevent entry and release of infectious virus genomes into host cells, likely making this responsible for the virucidal activity observed herein. How Rosin soap might disrupt the envelope is unknown, but Rosin soaps likely act as surfactants and further studies are required to determine this. Precisely how Rosin soap impacts the viral envelope is not known at this stage. Rosin soap is a mix of products, and that it would be useful to look at the individual compounds – both in terms of the resin acids and the carboxylic acids. However, there is limited commercial availability of these, and they also have limited solubility in pure solution. As has been done for other virucidal products (Fletcher et al., 2020).

Unfortunately, due to the cytotoxic nature of Rosin soap at high concentrations (from 0.25% to ~0.0025%) in our in vitro cell line cell culture conditions, we were not able to completely negate this background toxicity in our virus infectivity assays (which rely upon cellular integrity) even following purification of our virus/soap mixes by filtration. However, our data suggest that it is highly likely that Rosin soap inactivates the vast majority of infectious particles in a given prep. Using IAV, which grows to very high titres, we were able to demonstrate nearly complete inactivation. It is worth noting that this level of virus titre used in these experiments is higher than likely present in most ‘real world’ scenarios/environments (Boone and Gerba 2007). Despite our observation of toxicity in cell culture conditions, Rosin salves have been found to be safe and effective in wound care (Jokinen and Sipponen 2016).

The viricidal activity of Rosin soap when viruses are dried onto surfaces is an area that needs further research as viruses such as SARS-CoV-2 persist on surfaces and are a source of infection transmission (Kampf et al., 2020). This would determine if Rosin soap can be formulated into products that could be used as a commercial surface disinfectant for premises including hospitals. A wider variety of viruses could also be examined to determine if Rosin soap exhibits the same viricidal activity against most or all enveloped viruses such as SARS-CoV-2. Rosin soap did not inhibit the non-enveloped virus, EMCV. Other non-enveloped viruses, such as rhinoviruses or noroviruses could be examined to determine if it is only EMCV that rosin soap does not inhibit.

In conclusion, we demonstrate the virucidal activity of rosin soap against multiple pathogenic human enveloped viruses.

## Acknowledgements

The authors wish to thank Dr Ed Hutchinson (CVR), and Prof Ultan Power (QUB) for access to IAV and RSV, respectively. This work was funded by support from Hankkija Oy and Forchem Oy.

